# The effects of temperature on nestling growth in a songbird depend on developmental and social context

**DOI:** 10.1101/2025.10.05.680579

**Authors:** Sage A. Madden, Rebecca J. Safran, Gail L. Patricelli, Sara R. Garcia, Zachary M. Laubach

## Abstract

Climate change can adversely impact animals, especially those that cannot independently thermoregulate or avoid exposure. Cold, hot, and variable temperatures may impede nestling songbird growth due to increased thermoregulatory costs and reduced food delivery by parents. At a broad scale, temperature effects on nestling growth vary across climatic zones, but how temperature effects vary with early-life developmental constraints imposed by the timing of thermoregulatory development, competition with siblings, and the amount of parental care has received less attention. We investigated whether the effect of temperature on the mass of wild barn swallow (*Hirundo rustica erythrogaster*) nestlings (n = 113 nestlings, 31 nests) in Boulder County, CO depends on timing of exposure during development, relative size within the brood, or level of parental feeding. We found that the effect of minimum temperature differed in early and late development (before versus after putative development of thermoregulatory independence) and may be less pronounced for nestlings that received more parental feeding. We did not find evidence that the smallest nestling in the brood differs from other nestlings in vulnerability to temperature. These findings indicate the existence of fine-scale heterogeneity in which the effects of temperature on nestling development are sensitive to metabolic constraints and early-life social environment, providing key insight into the factors that may ameliorate or exacerbate climate impacts on individual birds.

## Introduction

Rising and increasingly variable temperatures due to climate change [1] can adversely impact animals, especially those that cannot independently thermoregulate or avoid exposure. At risk individuals include young animals lacking the mechanisms required to regulate exposure and cope with temperature extremes [2]. Exposure to excessively hot or cold temperatures is a form of developmental stress that negatively impacts not only the immediate survival of young animals in many species, but also the development of adult phenotypes [3,4]. Songbird nestlings may be particularly vulnerable to effects of extreme and variable temperatures because they are confined to a nest, ectothermic and mostly featherless early in development [5,6], and entirely dependent on parental care [7,8]. As a result, exposure to extreme or variable temperatures may impede nestling growth due to thermoregulatory costs and reduced food delivery by parent birds [9,10]. Although previous studies have revealed temperature effects on nestling growth across climate zones [9,10], we know little about how these effects may vary with localized early-life developmental constraints imposed by the timing of thermoregulatory development, competition with brood mates, and the amount of parental care. This knowledge is key for understanding heterogeneity in susceptibility of nestling birds to climate impacts and the factors that may ameliorate or exacerbate these impacts.

The effects of temperature on nestling development may differ depending on the timing of exposure during development [11–13]. Young nestlings initially lack the ability to thermoregulate independently and are mostly featherless [8,14], and body temperatures outside of a narrow range may disrupt growth [5,6,11,13]. The ability to thermoregulate develops partway through the nestling period [6,14], and extreme temperatures may require expending energy on thermoregulation and/or entering a state of hyperthermia, leading to increased stress and decreased growth [6,15,16]. The development of thermoregulatory independence partway through the nestling period suggests that effects of temperature may differ across nestling ontogeny. However, most studies have focused on the nestling period as a whole (but see [11–13,17]), leaving unanswered whether the effects of temperature differ before versus after development of thermoregulatory independence.

A nestling’s social environment also may mitigate or exacerbate the effects of temperature on their development [18,19]. Songbird nestlings often hatch asynchronously, resulting in size asymmetry that is maintained across development [20]. This asymmetry may affect access to resources—older nestlings outcompete younger nestlings for food, and younger nestlings often grow more slowly and even die from starvation [20–22]. Heterogeneity in access to resources within a brood may then drive differences in susceptibility to temperature. Additionally, body size is a critical determinant of thermoregulatory ability [6,23], but most studies have investigated the relationship between size and thermoregulation among rather than within species [24,25]. As such, investigating whether the size of a nestling relative to its brood mates influences responses to temperature can help elucidate how early-life social environment shapes nestling responses to ecological conditions [21,26–28].

Parental care is another aspect of the social environment that may shape nestling responses to temperature [18,19,29]. Nestlings are entirely dependent on food provisioned by parents to provide energy for growth and thermoregulation. Nestlings that receive more parental care may be less susceptible to the negative effects of temperature because they have more energetic reserves for thermoregulation and growth. Indeed, research on burying beetles (*Nicrophorus vespilloides*) indicates that higher levels of parental care can buffer young from effects of temperature variation [30]. Similarly, studies of cooperatively breeding bird species suggest that the presence of helpers at the nest may mitigate negative impacts of weather on nestlings [18] (but see [29]). Therefore, it is plausible that parental care influences nestling vulnerability to extreme temperatures in songbirds.

In this study, we investigated the effects of temperature at different stages across nestling ontogeny in wild barn swallows, and we explored whether nestling size relative to brood mates or level of parental feeding modify the effects of temperature on nestling growth. We aimed to address the following three questions (Fig 1): 1) Does the effect of temperature on nestling mass differ when the exposure is assessed during early versus late development? 2) Does the effect of temperature on nestling mass depend on whether a nestling is the smallest in the brood? 3) Does the effect of temperature on nestling mass depend on parental care? For question one, we hypothesized that exposure to extreme and variable temperatures would impose greater metabolic costs on nestlings before thermoregulatory independence than after, leading to stronger negative effects of temperature on mass early in development than late. For questions two and three, we hypothesized that advantageous social environments (e.g., larger size within the brood hierarchy, higher levels of parental feeding) would ameliorate some of the negative impacts of extreme and variable temperatures on nestling mass due to the nestlings’ greater access to resources. Directed acyclic graphs [31] displaying hypothesized relationships investigated for each question are provided in S1-S3 Fig.

**Fig 1.**
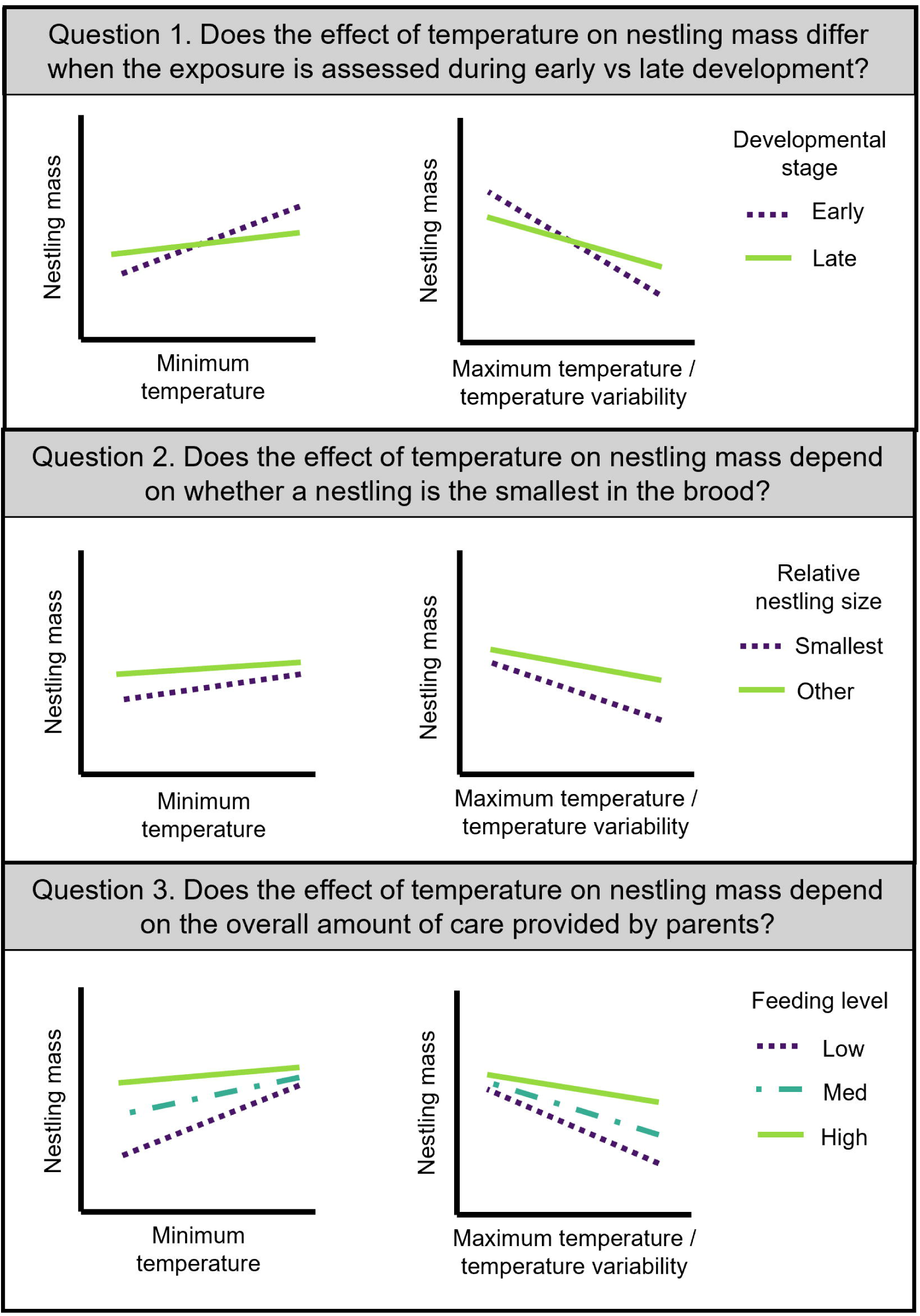
Research questions and predicted results for our hypotheses about temperature effects on wild barn swallow nestling mass before fledging. We expected that lower minimum temperature, higher maximum temperature, and greater temperature variability (interquartile range) would be associated with lower nestling mass. Feeding level refers to the average nest-level feeding rate across conditions.

## Methods

### Study system

We monitored nesting attempts from wild barn swallows during the summer of 2021 at seven breeding sites in Boulder County, CO. These sites are monitored as part of a long-term study of barn swallows and are occupied by colonies ranging in size from one to 50 breeding pairs. Barn swallows build open mud cup nests and lay clutches of three to five eggs, which hatch asynchronously over approximately 1-2 days [32]. Hatching asynchrony results in a size hierarchy that persists over the course of development [33]. Hatch order is not associated with offspring sex and there are no morphological differences between different sexes [34]. In this socially monogamous species, both parents in a social pair provide care, including food provisioning, until nestlings fledge (>18 days post hatch) [32].

### Study design overview

We collected environmental, demographic and behavioral data at 31 first brood nests from May through July 2021, and we measured morphometric and physiological traits on 113 nestlings across development (Fig 2). Nests were checked every three to five days to track nestling phenology and fate. During the first check after nestlings hatched, we estimated their ages (hatch day = day zero) based on feather emergence and other reliable developmental characteristics, such as wetness (on hatch day) and ability to raise their head [35]. We took morphological measurements of nestlings and conducted one-hour observations of parental care behavior at three time points: days three to four, days eight to nine, and days 11-13. We monitored temperature near the nest from hatch through last nestling measures.

**Fig 2.**
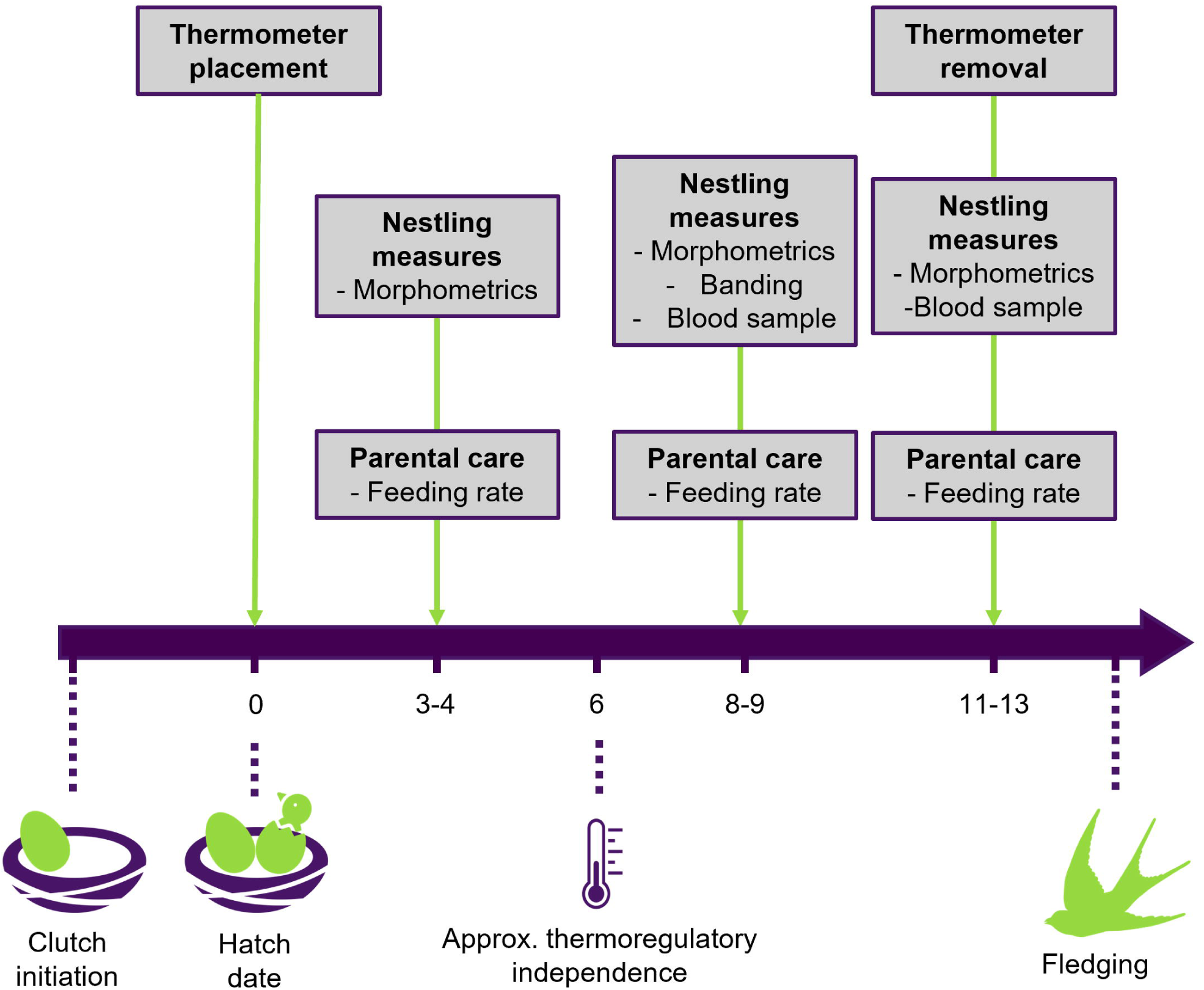
An overview of the study design indicating the types of data collected across the nestling period at three time points in wild barn swallows in Boulder County, CO. We expect thermoregulatory independence to develop at ∼six days post-hatch in barn swallows (see “Near-nest temperatures”).

### Nestling measures

We collected standard morphometric measurements for each nestling at three time points ([36]; Fig 2). We measured mass to the nearest 100th of a gram using a digital scale and right-wing length to the nearest half millimeter as the distance from the carpal joint to the distal end of the second phalanx or primary wing feather, whichever was longer. On days eight to nine, we placed a USGS metal band on the tibiotarsus of each nestling, allowing us to track individuals through development. On days eight to nine and 11-13, we also collected two small blood samples from each nestling as part of a separate study [36].

Using day eight to nine measures (the earliest age at which we could begin tracking individuals), we created a categorical variable of relative nestling size, where the smallest nestling in each nest based on right-wing length was classified as ‘min’ (reference group) and the remaining nestlings were classified as ‘other’. Our classification was based on right wing length, rather than mass, because wing length measures skeletal size and may not be as strongly influenced by short-term fluctuations as mass [37–39]. Because we were unable to track individuals until days eight to nine, we could not determine the exact age of individual nestlings. Therefore, we are unable to tease apart the effects of age and mass in the relationship between relative nestling size and vulnerability to adverse temperatures.

### Parental care observations

We conducted ∼one-hour focal observations of each nest at three time points (see details in [36]; Fig 2). Some nests are missing data for one or more time points due to logistical issues. The start of each observation was preceded by collection of morphological measurements and blood samples from nestlings (see Nestling Measures). After these measures, observers watched the nest from a blind, and the birds were given ∼15 minutes to habituate to our presence before the observation commenced (following [36,40]). Observations began between 5:45 and 8:00 am. During each observation, the observer logged parental care behavior in real-time on iPads or iPhones (Apple Inc.) using “Animal Behavior Pro” [41]. Behaviors recorded included feeding visits, where the parent delivers food to a nestling (see full ethogram in S1 Table). Given biparental care in this species, we recorded the total number of behaviors for both social parents.

When an observer could not be present, we instead recorded nests using GoPro cameras, and videos were scored by the same observers who conducted the in-person observations. For these camera observations, we set up camera mounts the day before recording to allow birds to habituate, and we waited for ∼15 minutes after the cameras were started to begin scoring behavior. Parental care behaviors are consistent when scored by different observers and between in-person and camera observations [40].

We created an index of the level of parental care by summarizing feeding rates from both parents across all stages of nestling development. Specifically, this index was created using a generalized linear mixed effects model, in which total feeding count by both parents measured at each developmental stage was the outcome and nest ID was a random intercept. We compared model fit for several error distributions and link functions using diagnostic plots from the package ‘DHARMa,’ version 0.4.6 [42], and we selected a negative binomial distribution and log link for our final model. The model included an offset for total observation time and several covariates that could influence parental feeding behaviors: number of days since the first nestling hatched, number of nestlings in the nest, the median temperature near the nest during the observation period, and the duration of time between removing nestlings from the nest and the start of the observation. From this model, we extracted the best linear unbiased predictors (BLUPs) and created a three-level variable of parental feeding based on nest-level tertiles of average feeding rates summarized across development (following [36]).

Using a mixed effects model in this way allowed us to summarize parental care across developmental stages while controlling for environmental conditions and nest-level variables that may influence parental care. Moreover, this method is an efficient use of our data because it allows for missing data via shrinkage of nest-level estimates toward the global sample mean. Because parental feeding is an effect modifier in our models, we created a three-level categorical variable for stratified analyses by classifying BLUPs as ‘low’ (lowest third), ‘med’ (middle third), or ‘high’ (highest third) parental feeding. The approach of carrying BLUPs forward from one model to another has been criticized due to its failure to propagate uncertainty [43]. However, because summarizing parental feeding using BLUPs should not substantively shift nests from one parental feeding level to another, this approach is justified in our case given the advantages described above.

### Near-nest temperatures

We hung thermometers (Govee) in small mesh bags adjacent (10-28 cm) to each open cup nest and logged near-nest temperature and humidity every 15 minutes throughout the data collection period (Fig 2). At one nest, we are missing two days of temperature measurements at the end of the nestling period. We calculated the minimum temperature, maximum temperature, and temperature variability (defined as the interquartile range) at each nest from day zero (hatch) to the end of day five (early development) and from day six until the last measure of nestling mass on day 11-13 (late development). These temperature summaries correspond with before and after the putative development of thermoregulatory independence, respectively. We chose these time periods based on previous research in swallows and other songbirds [14,44–46], which found that nestling swallows begin to develop the ability to thermoregulate independently around four to five days post-hatch, and the age of the effective homeothermy (defined here as ability to maintain relatively constant body temperature under natural conditions—in the nest with brood mates) is likely around six days of age. Finally, we calculated the minimum temperature, maximum temperature, and temperature variability at each nest over the entire nestling period, from hatch day to the last measure of nestling mass.

### Ethics statement

All research on wild barn swallows was approved by the Rebecca Safran’s Institutional Animal Care and Use Committee protocol (permit no. 1303.02), and all research was performed in accordance with relevant guidelines and regulations. In addition, this work was conducted under the Bird Banding Lab, master permit bander ID 23505. Capture using mist nests in Boulder, Weld and Jefferson Counties, Colorado, was approved by the Colorado Parks and Wildlife issued under permit license number 21TRb2005. Some swallow nests were located on private property and were monitored with permission from the landowners.

### Statistical analysis

We conducted three sets of formal analyses aligning with our questions. Across all questions, we investigated: 1) minimum temperature, which captures exposure to extreme cold, 2) maximum temperature, which captures exposure to extreme heat, and 3) temperature variability, which captures exposure to fluctuations between cold and hot [9,10]. Given our modest sample size, we balanced our inference based on the consistency, direction, and magnitude of effect estimates alongside of conventional significance thresholds. We focused our reporting and interpretation of results on estimates of effect sizes and 95% confidence intervals, providing information on the strength and likely error of our effect estimates [47], as well as p-values interpreted using the language of evidence [48]. P-values less than 0.05 were interpreted as providing moderate to strong evidence for an effect, p-values between 0.05 and 0.10 as providing marginal evidence, and p-values greater than 0.10 as providing little to no evidence [48].

Across all analyses we report both unadjusted associations and adjusted associations from multiple variable regression models; Examining both unadjusted and adjusted associations allowed us to see if the estimate of interest changes after controlling for other variables. For adjusted models, we included hatch date and number of nestlings in a nest as potential confounders. For all models, we z-score standardized numeric explanatory variables and covariates to aid in model fit and comparison of effect sizes. All models included a random intercept for nest ID to account for non-independence of measures taken at the same nest. We originally included a random effect for site ID to account for non-independence of measures taken at the same site, but this effect was removed because it explained zero variance in some models. Removal of site ID from models in which it explained some variance did not substantially affect the direction, magnitude, or significance of effects.

Model assumptions were checked using diagnostic plots. Specifically, we checked whether each model met the assumptions of a Gaussian error distribution with homogenous variance by examining plots of residuals versus fitted values, a histogram of the residuals, a normal quantile-quantile plot, and Cook’s distance and standardized residual influence plots, using the package ‘car’, version 3.1.3 [49]. For some models, we detected influential outliers, which we defined as having a Cook’s distance greater than one or a standardized residual value greater than three. When influential outliers were present, we conducted sensitivity analyses, where we ran the model with and without the outlier(s) and compared the results.

For question one, we separately ran two sets of linear mixed models for each of our three temperature variables (minimum, maximum, and variability) (S1 Fig): 1) In the first set of three models, each temperature variable collected in early development (from hatch to the end of day five) was included as the explanatory variable of interest and nestling mass on days 11-13 was the outcome; 2) the second set of models were the same except that temperature variables were quantified in late development (from day six to last nestling measures on days 11-13). We conducted a sensitivity analysis to assess the robustness of our results to differences in the selected cut-off for development of thermoregulatory independence. Specifically, we ran the same set of models described above, except using cut-off ages of five days post-hatch and seven days post-hatch, rather than six days post-hatch, for development of thermoregulatory independence.

For questions two and three, we tested for effect modification of temperature effects on nestling mass by relative nestling size and level of parental feeding, respectively. Effect modification refers to a situation in which the effect of X on Y depends on a third variable, the effect modifier [31]. In our analysis, relative nestling size (question two) and level of parental feeding (question three) were treated as effect modifiers. To determine whether effect modification was present, we ran analyses stratified by multiple levels of relative nestling size or parental feeding. For question two, we ran models of the effect of each temperature variable on mass stratified by nestling size, ‘min’ = the smallest nestling in the nest and ‘other’ = all other nestlings. For question three, we modeled the effect of temperature on mass stratified by parental feeding, which included three levels, ‘low’ = parental feeding below the ∼33^rd^ percentile, ‘med’ = parental feeding between the ∼33^rd^ and ∼67^th^ percentile, and ‘high’ = parental feeding above the ∼67^th^ percentile. S4 Fig displays the feeding rate BLUPs, grouped into three strata. The results from these stratified models provide insight into the magnitude and directionality of the effect of temperature on nestling mass for different relative nestling sizes or levels of parental feeding.

All statistical analyses were carried out using R version 4.4.1 [50]. The ‘tidyverse’ package, version 2.0.0 [51] was used for data cleaning and organization. The ‘lme4’ package, version 1.1.35.5 [52] was used to fit models. Bootstrapped confidence intervals were obtained for model coefficients using the package ‘boot’, version 1.3.30 [53,54]. P-values for two-tailed t-tests for each coefficient were obtained using Satterthwaite degrees of freedom method, via the package ‘lmerTest’ version 3.1.3 [55]. Goodness of fit was assessed using conditional and marginal R-squared values, calculated using the package ‘MuMIn’, version 1.48.4 [56]. Results were visualized using the package ‘ggplot2’ version 3.5.1 [57], ‘ggeffects’ version 1.7.2 [58], and ‘ggpubr’ version 0.6.0 [59].

## Results

### Background characteristics

Of the 113 nestlings in this study, 108 survived until days eight to nine, and 106 survived until days 11-13. Eggs hatched asynchronously over up to two days—the mean difference in age between the first and last hatched nestling was 1.03 days (range = 0-2 days). For the near-nest temperature measurements averaged across the nestling period, the mean ± SD was 13.10 ± 1.80 degrees C for minimum temperature, 36.90 ± 2.96 degrees C for maximum temperature, and 9.26 ± 2.43 degrees C for temperature variability. We categorized nestlings based on within nest relative size on days eight to nine, measured in terms of right-wing length. At this time, the mean ± SD right wing length was 26.8 ± 5.55 mm for the smallest (‘min’) nestlings and 32.4 ± 4.26 mm for ‘other’ nestlings. The mean ± SD days eight to nine mass was 11.60 ± 2.61 grams for the smallest (‘min’) nestlings and 13.60 ± 2.30 grams for ‘other’ nestlings. On days 11-13, the timepoint at which the outcome was measured, the mean ± SD wing length was 49.20 ± 8.09 mm for the smallest (‘min’) nestlings and 54.40 ± 4.76 mm for ‘other’ nestlings. The mean ± SD days 11-13 mass was 17.20 ± 2.75 grams for the smallest (‘min’) nestlings and 17.70 ± 1.91 grams for ‘other’ nestlings. For measures of parental care, the mean ± SD total feeding rate (counts/hour) was 12.21 ± 6.92 on days three to four, 12.73 ± 8.08 on days eight to nine, and 21.62 ± 12.94 on days 11-13. Feeding rate and proportion of time spent brooding during the observation were negatively correlated (Spearman Rank Correlation: rho = −0.49, p < 0.001). Additional background characteristics for the study population are provided in S2 Table. S5 Fig displays daily minimum temperature, maximum temperature, and temperature variability across the nestling period for each nest.

### Question 1. Does the effect of temperature on nestling mass differ when the exposure is assessed during early versus late development?

Higher minimum temperatures in early development (β = 1.16 g per 1 SD [95% CI: 0.32, 2.00], P = 0.01) but not late development (β = 0.29 g per 1 SD [95% CI: −0.45, 1.06], P = 0.46) were associated with higher nestling mass at last measure (S3 Table; Fig 3a). Higher maximum temperatures in early (β = −0.66 g per 1 SD [95% CI: −1.32, −0.04], P = 0.05) and late (β = −1.13 g per 1 SD [95% CI: −1.74, −0.52], 0.001) development were associated with lower nestling mass (S3 Table; Fig 3b). Similarly, greater temperature variability in early (β = - 1.41 g per 1 SD [95% CI: −2.07, −0.77], P = 0.0003) and late (β = −1.33 g per 1 SD [95% CI: −1.95, −0.70], P = 0.0003) development was associated with lower nestling mass (S3 Table; Fig 3c).

**Fig 3.**
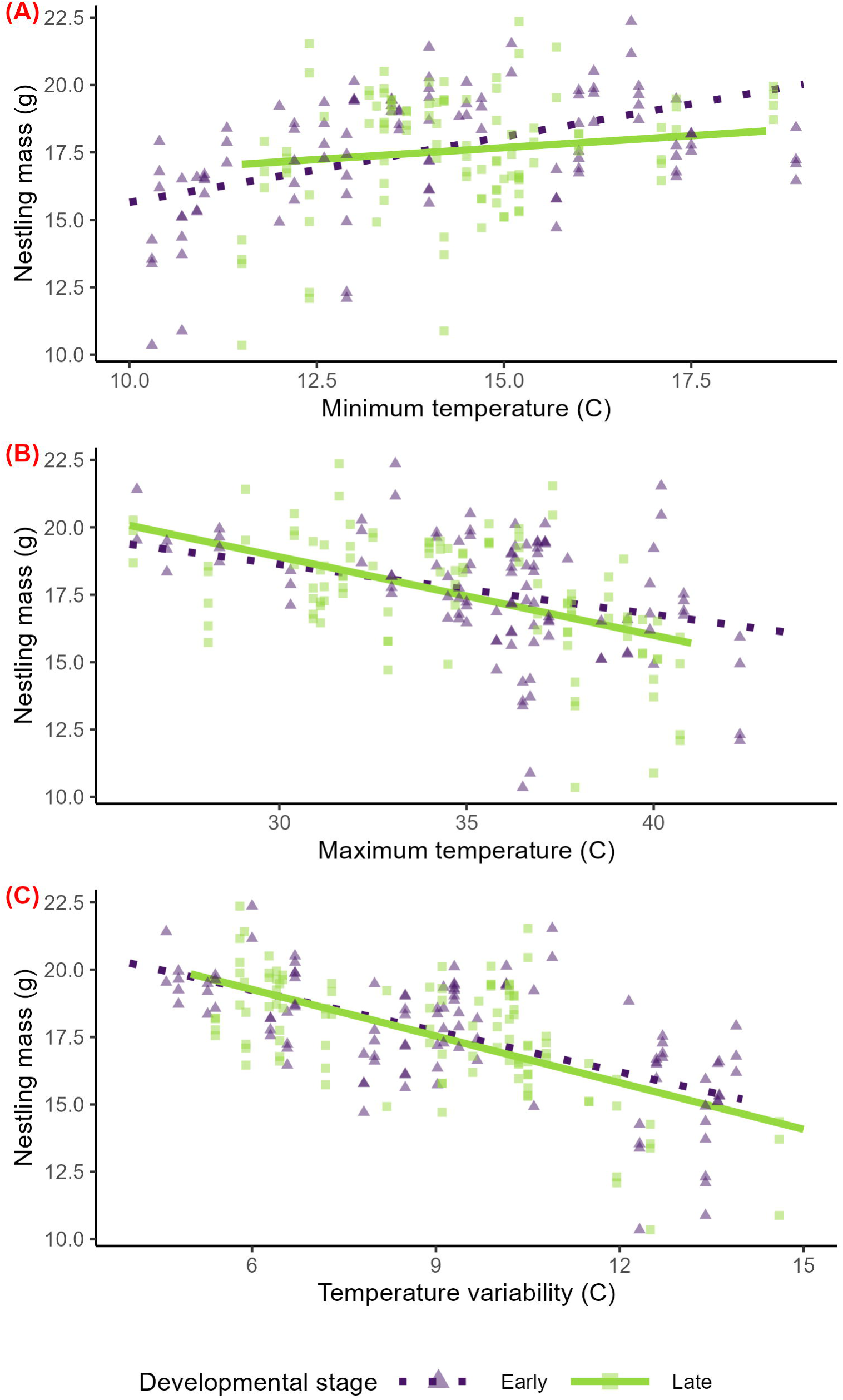
Temperature effects in early and late development on the mass of 11–13-day old wild barn swallow nestlings in Boulder County, CO. Predicted relationships of minimum temperature (a), maximum temperature (b), and temperature variability (c) and nestling mass from separate linear mixed models for each developmental stage—in early (‘Early’) development, from hatch to end of day five days post-hatch, and late (‘Late’) development, from six days post-hatch to final measures at 11-13 days post-hatch. Model predictions are displayed as lines and raw data are displayed as points. Colors, line types, and shapes correspond to developmental stage.

The estimates of interest did not change substantively when influential outliers were excluded from analyses or when models were run using five days or seven days post-hatch, rather than six days post-hatch, as the cut-off for age for early versus late development.

### Question 2. Does the effect of temperature on nestling mass depend on whether a nestling is the smallest in the brood?

In stratified analyses, higher minimum temperatures across the nestling period corresponded with higher nestling mass for both the smallest nestlings in the nest (β = 1.75 g per 1 SD [95% CI: 0.43, 3.14], P = 0.02; S4 Table; Fig 4a) and for other nestlings (β = 1.30 g per 1 SD [95% CI: 0.40, 2.21], P = 0.01; S4 Table; Fig 4a). Similarly, higher maximum temperatures and greater temperature variability across the nestling period were associated with lower nestling mass for the smallest nestlings in the nest (maximum: β = −1.42 g per 1 SD [95% CI: −2.48, −0.40], P = 0.01; variability: β = −1.95 g per 1 SD [95% CI: −2.98, −0.92], P = 0.001; S4 Table; Fig 4b-4c) and for other nestlings (maximum: β = −0.85 g per 1 SD [95% CI:-1.47, −0.27], P = 0.01; variability: β = - 1.50 g per 1 SD [95% CI: −2.15, −0.89], P = 0.0001; S4 Table; Fig 4b-4c).

**Fig 4.**
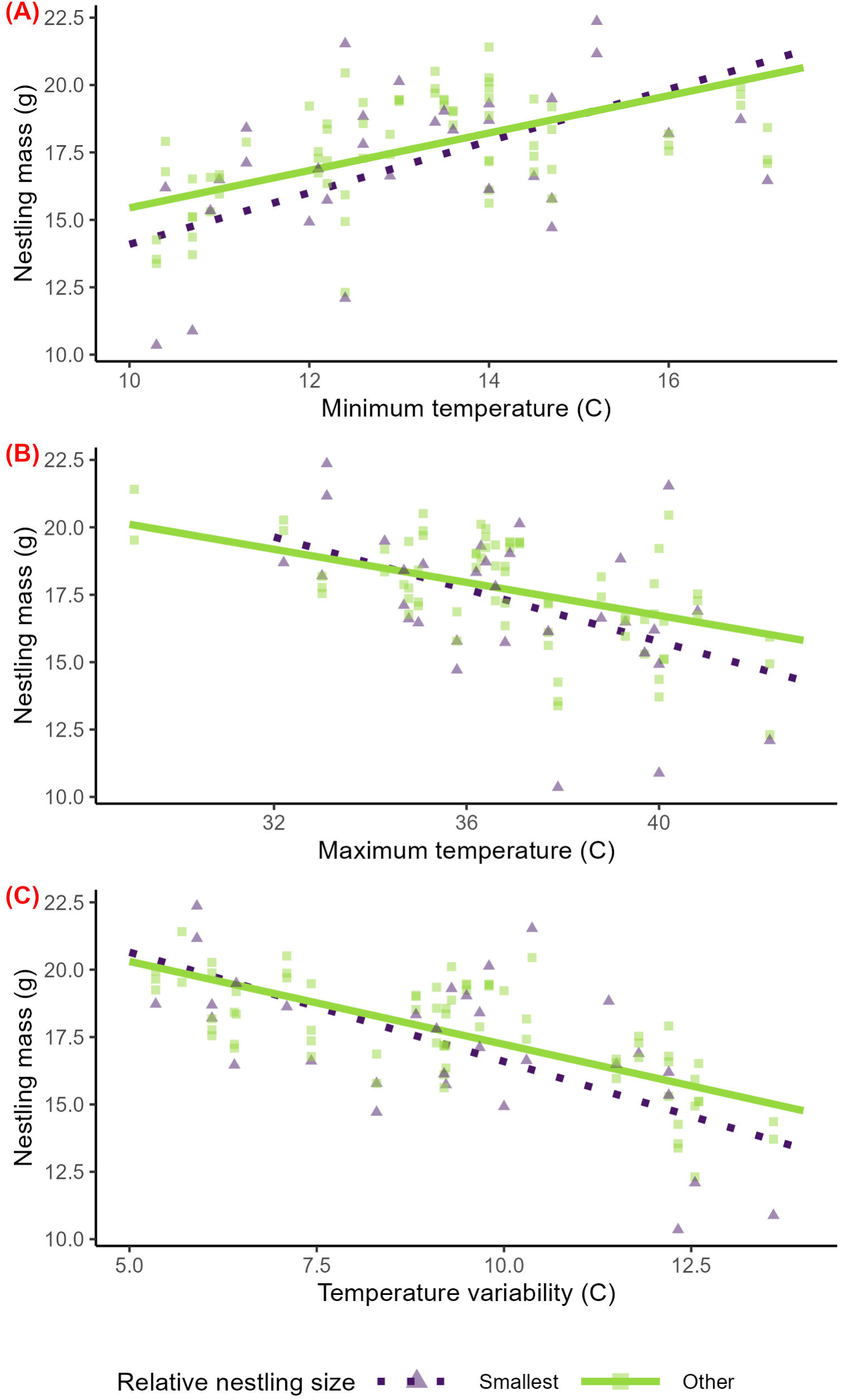
Temperature effects on the mass of 11-13-day old wild barn swallow nestlings in Boulder County, CO that are the smallest in their brood versus other nestlings. Predicted relationships of minimum temperature (a), maximum temperature (b), and temperature variability (interquartile range) (c) across the nestling period and nestling mass from linear mixed models, stratified by relative nestling size—the smallest nestling (‘Smallest’) and all other nestlings (‘Other’)—at days eight to nine post-hatch. Model predictions are displayed as lines and raw data are displayed as points. Colors, line types, and shapes correspond to relative nestling size. Because relative nestling size was determined at days eight to nine and some nestlings did not survive to final measures on days 11-13, for some nests, only ‘Smallest’ or ‘Other’ nestling points are present.

### Question 3. Does the effect of temperature on nestling mass depend on the overall amount of care provided by parents?

In stratified analyses, higher minimum temperatures across the nestling period were associated with higher nestling mass among nests that received low parental feeding, but the confidence interval overlapped zero (β = 1.73 g per 1 SD [95% CI: −0.11, 3.53], P = 0.11; S5 Table; Fig 5a). Higher minimum temperatures were also associated with higher nestling mass among nests that received medium parental feeding (β = 2.36 g per 1 SD [95% CI: 0.68, 3.92], P = 0.03; S5 Table; Fig 5a). In contrast, we observed no clear association of minimum temperatures with nestling mass among nests that received high parental feeding (β = −0.53 g per 1 SD [95% CI: −2.05, 0.94], P = 0.53; S5 Table; Fig 5a).

**Fig 5.**
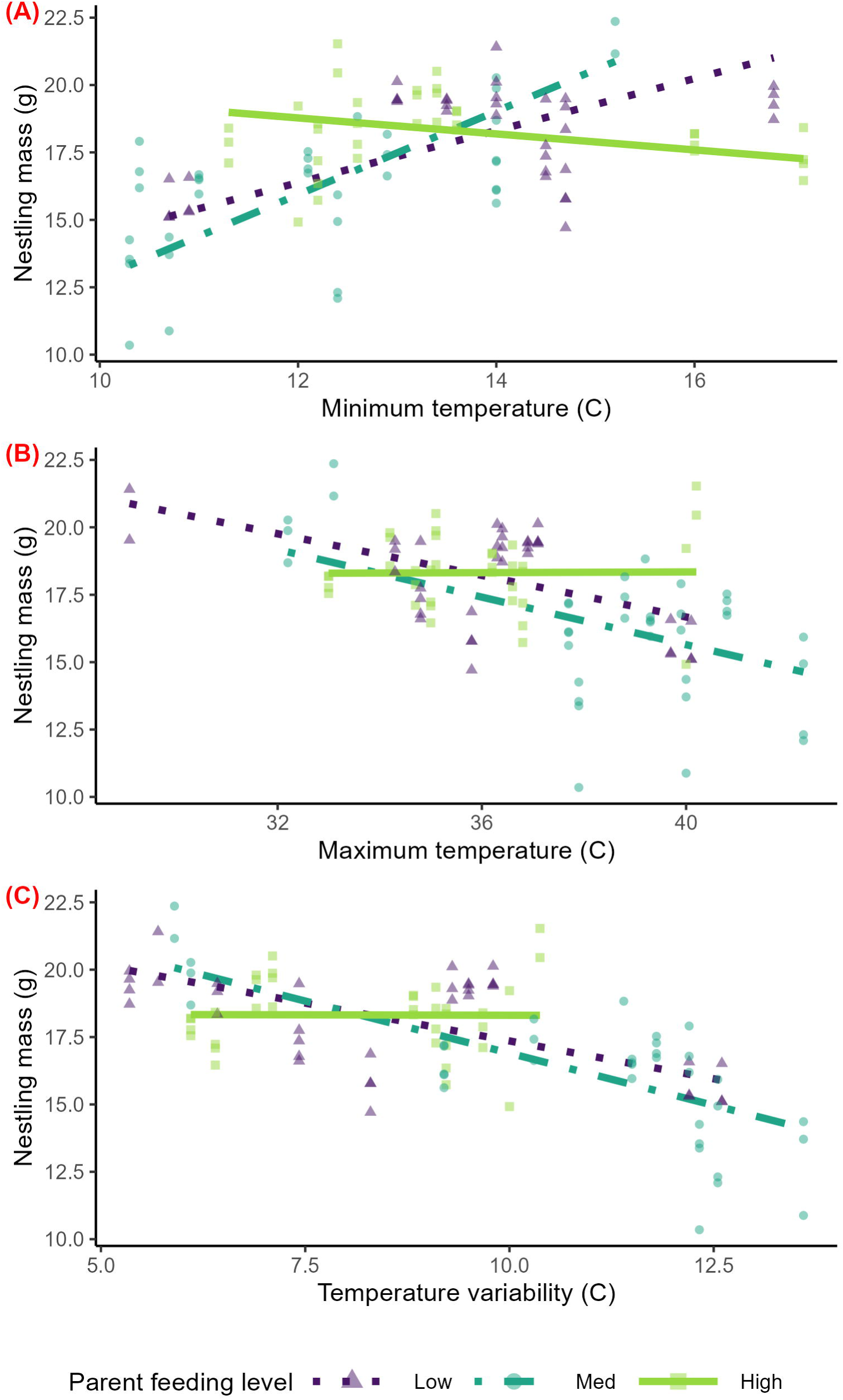
Temperature effects on nestling mass of 11–13-day old wild barn swallows in Boulder County, CO at nests receiving low, medium, or high parental feeding. Predicted relationships of minimum temperature (a), maximum temperature (b), and temperature variability (interquartile range) (c) across the nestling period and nestling mass from linear mixed models, stratified by level of parent feeding—low (‘Low), medium (‘Med’), and high (‘High’). Model predictions are displayed as lines and raw data are displayed as points. Colors, line types, and shapes correspond to level of parent feeding.

Similarly, higher maximum temperatures and greater temperature variability across the nestling period were associated with lower nestling mass among nests that received low parental feeding, but confidence intervals included zero (maximum: β = −1.08 g per 1 SD [95% CI: - 2.44, 0.34], P = 0.16; variability: β = −1.35 g per 1 SD [95% CI: −2.77, 0.14], P = 0.11; S5 Table; Fig 5b-c). Higher maximum temperatures and greater temperature variability were also associated with lower nestling mass among nests that received medium parental feeding (maximum: β = −1.22 g per 1 SD [95% CI: −2.34, −0.04], P = 0.07; variability: β = −1.80 g per 1 SD [95% CI: −2.78, −0.75], P = 0.01; S5 Table; Fig 5b-c). In contrast, we observed no clear association of maximum temperature or temperature variability with nestling mass among nests that received high parental feeding (maximum: β = 0.01 g per 1 SD [95% CI: −0.91, 0.92], P = 0.98; variability: β = −0.01 g per 1 SD [95% CI: −1.25, 1.28], P = 0.99; S5 Table; Fig 5b-c).

The magnitude, direction, and precision of the estimates of interest did not substantively change when influential outliers were excluded from analyses.

## Discussion

In this study of 113 wild barn swallow nestlings in 31 nests, we investigated whether the effects of minimum temperature, maximum temperature, and temperature variability on nestling mass differ according to the timing of exposure across nestling ontogeny and two features of the nestlings’ social environment: the nestlings’ size relative to brood mates and the level of parental feeding the nest received. We found that exposure to lower minimum temperatures, higher maximum temperatures, and greater temperature variability were associated with lower nestling mass. Additionally, we found some evidence for heterogenous effects of temperature on nestlings across developmental stages and social environments. When looking at the effects of exposure to temperatures in early versus late development (before versus after putative development of thermoregulatory independence), we found different relationships of minimum temperature with nestling mass. Surprisingly, the association of temperature with nestling growth did not vary with relative nestling size, suggesting that sibling size dynamics may not have a large impact on nestling vulnerability to adverse temperatures. Finally, we found that the relationship between temperature and nestling mass observed at medium levels of parental feeding were absent from high parental feeding nests. Our work adds to relatively limited knowledge of how temperature effects on nestlings vary with developmental constraints imposed by timing of thermoregulatory development and aspects of early-life social environment, which is critical to understanding factors that may ameliorate or exacerbate climate change impacts on birds.

### Minimum temperature, maximum temperature, and temperature variability have differential effects on nestling mass

Our results align with literature reporting that exposure to cold, hot, and variable temperatures may all lead to reduced nestling growth. We found that lower minimum temperatures were generally associated with lower nestling mass, consistent with previous studies finding that exposure to cold temperatures leads to reduced nestling growth [11,60,61]. On the other hand, higher maximum temperatures were generally associated with lower nestling mass in our population as in other studies [62–64]. Greater temperature variability was generally associated with lower nestling mass, consistent with evidence that nestling growth is reduced when exposed to more variable than stable temperatures [65].

Notably, some studies report no effect of extreme or variable temperatures on nestling growth [66–68]. Ecological and social factors, such as habitat, diet, and nest type, may shape susceptibility to negative effects of temperature across species and populations [18,69]. For example, insectivorous species may be more sensitive to extreme temperatures than granivorous species because insect availability is more strongly influenced by weather conditions [70,71]. Additionally, open-cup nesting species may be more sensitive to extreme temperatures than cavity-nesting species because open-cup nests may be less thermally buffered than cavity nests [69]. Given that barn swallows are insectivorous and build open cup nests, it is perhaps unsurprising that nestling mass is negatively affected by cold, hot, and variable temperatures in this species.

### Effects of temperature on nestling mass may differ across nestling ontogeny

Consistent with our hypothesis, we found that lower minimum temperatures in early but not late development--*before but not after* putative development of thermoregulatory independence at six days post-hatch—were associated with lower nestling mass (Fig 3a). Nestlings may be especially sensitive to cold temperatures before thermoregulatory independence because colder ambient temperatures result in colder body temperatures at times when the parents are away from the nest, and colder temperatures slow metabolic processes in young nestlings [72]. In older nestlings, improved thermoregulation allows for a more constant body temperature and metabolic rate increases when ambient temperatures are outside the thermal neutral zone [6]. Additionally, due to their higher surface area to volume ratio and lack of feathers, young nestlings may dissipate heat more quickly than older nestlings, which may be a disadvantage under cold conditions [6,8,68].

Contrary to our hypothesis, higher maximum temperatures and greater temperature variability in early and late development—*before and after* putative development of thermoregulatory independence*—*were associated with lower nestling mass (Fig 3b-c). This suggests that nestlings may be sensitive to heat and environmental instability regardless of thermoregulatory status, though previous work indicates that the nature of thermal challenges may vary across development. Early in development, nestlings may be susceptible to negative impacts of extreme heat because they may gain heat more quickly than older nestlings due to their high surface area to volume ratio and lack of feathers [6,8,68]. Late in development, after development of thermoregulatory independence, nestlings may be sensitive to hot temperatures because evaporative cooling is energetically costly and may quickly lead to dehydration [64,73]. In addition, nestlings may have limited ability to cool themselves [73], and this may be exacerbated in older nestlings by reduced heat dissipation due to higher surface area to volume ratio, feather development, and crowding in the nest (though as noted above, reduced heat transfer may be an advantage when ambient temperatures exceed body temperature). Finally, older nestlings have higher energetic requirements, which are likely near or in excess of the limits of the food parents can provision [74,75], and they may be more strongly affected by changes in food abundance or parental care behavior than younger nestlings [11].

### Little evidence that effects of temperature on nestling mass differ by relative size

Contrary to our hypothesis, we did not find evidence for effect modification by relative nestling size on the relationship between temperature and nestling mass—lower minimum temperatures, higher maximum temperatures, and greater temperature variability were associated with lower nestling mass regardless of relative nestling size (Fig 4).

Our findings contrast with previous work suggesting that animals with a smaller body size are more vulnerable to extreme temperatures because they are less able to regulate their body temperature—they dissipate heat more quickly when ambient temperature is below body temperature, and they absorb heat more quickly when ambient temperature exceeds body temperature [6,23]. Consistent with this idea and in contrast to our findings, a study of tree swallows found that the smallest nestling in a brood had a lower chance of surviving harsh cold snaps than larger nestlings [26]. Similarly, in studies of black kites (*Milvus migrans*) and red-winged blackbirds (*Agelaius phoeniceus*), higher maximum and mean temperatures were linked to faster growth in the last hatched nestling, but not other nestlings [21,28].

Our findings are also surprising given that smaller nestlings may face a competitive disadvantage compared to their brood mates. Larger nestlings often outcompete younger nestlings for resources, which are crucial thermoregulation and other developmental processes like growth [20,21]. For example, in a study of blue tits (*Cyanistes caeruleus*), higher levels of rainfall had stronger positive impacts on the mass of early hatched nestlings than that of later hatched nestlings, which the authors suggested may be due to the greater ability of early hatched nestlings to secure food from parents than late hatched nestlings [27]. Additionally, competitive dynamics may shift under adverse conditions—extreme temperatures may exacerbate the competitive disadvantage of small nestlings [76]. Therefore, we might expect that larger nestlings that more efficiently thermoregulate or that can acquire more resources are likely to be less susceptible to extreme temperatures.

In contrast to this idea and findings previous studies, the lack of effect modification we observed may reflect that the sibling size hierarchy does not strongly affect nestling vulnerability to temperature in our study system. This may be because the differences in size among siblings were relatively small—at the timepoint when relative size was determined in our study, the mean ± SD right wing length was 26.8 ± 5.55 mm for the smallest (‘min’) nestlings and 32.4 ± 4.26 mm for ‘other’ nestlings.

### Effects of temperature on nestling mass may differ by level of parental feeding

In support of our hypothesis, we found some evidence for effect modification by parental feeding level for the association of temperature with nestling mass. Stratified models showed that lower minimum temperatures, higher maximum temperatures, and greater temperature variability were associated with lower nestling mass at nests receiving medium, but not high, levels of parental feeding (Fig 5). There was also a pattern of cold, hot, and variable temperatures being associated with lower nestling mass at nests receiving low levels of parental feeding, though the confidence intervals for these associations overlapped zero.

These findings suggest that high levels of parental feeding may mitigate some of the negative effects of adverse conditions on nestling development and are consistent with several studies in non-avian species and cooperatively breeding birds. As previously mentioned, maternal care in burying beetles reduced the adverse effects of cold temperatures on larvae.

Similarly, in sociable weavers (*Philetairus socius*), a cooperatively breeding bird species, having additional helpers at a nest counteracted some negative effects of adverse weather conditions (periods of low rainfall) on fledgling condition [18]. However, another study in the same species reported that having larger groups feeding nestlings did not mitigate the negative effects of adverse weather [29].

In addition to feeding, parental care may also involve other behaviors, like brooding, that influence nestlings’ responses to temperature exposure. For example, if parental feeding and brooding positively covary, as some studies have reported [77,78], then the effect of parental care on nestling mass maybe driven by active parental regulation of nest temperatures, in addition to or instead of access to higher energetic resources via food provisioning. However, in our study population (and other studies: [79,80]), feeding and brooding are negatively correlated, suggesting that positive covariation among these behaviors is not driving our observed differences in temperature effects across parental feeding levels.

Finally, it would be valuable for future work to tease apart the roles of average levels of parental care across development, as we have done, in addition to considering changes in specific bouts of parental care in response to temperature, in shaping effects of temperature on nestling development. In other words, parental care may act as a modifier of temperature effects on nestlings if these effects differ with average level of care, and as a mediator temperature effects if extreme temperatures cause reduced care, leading to reduced nestling growth. An increasing body of work documents the role of parental care as a mediator of temperature effects on nestlings—parent provisioning rates may decrease in hot or cold temperatures due to reduced availability of insect prey [11,81] and parents diverting more time and energy to thermoregulation, instead of parental care [82,83]. These changes in parental provisioning behavior during extreme weather conditions may then lead to reduced nestling survival and growth [11,63]. However, studies considering parental care as a modifier of temperature effects seem to be limited, and considering parental care as both an effect modifier and mediator warrants further investigation.

### Limitations and strengths

This study was not without limitations. First, our sample size may limit our ability to detect weak associations and differences in magnitude among strata, which may have led to some of our marginal and non-significant findings; further work with larger sample sizes is needed to provide stronger quantitative evidence for these patterns. Second, we modeled three temperature variables (minimum, maximum, and variability) separately due to limitations of our statistical power; further work with a larger sample size is needed to determine the relative strength of the effects of different temperature variables on nestlings or whether there are cumulative effects of different types of adverse temperature exposures (see discussion in [84,85]). Third, we examined differential effects of temperature across nestling ontogeny by categorizing the nestling period into two time periods, before and after putative development of thermoregulatory independence, based on previous studies of other species of swallows and songbirds [14,44–46]. Further studies are needed to understand how vulnerability to adverse temperatures changes during the gradual development of thermoregulatory independence [46,72]. Fourth, because we were unable to track individual nestlings from hatch, further work is needed to differentiate the effects of age and size in the relationship between relative nestling size and vulnerability to extreme and variable temperatures, such as by manipulating hatch asynchrony [86]. Additionally, degree of hatch asynchrony might be related to nest temperatures during incubation, but we were unable to assess this because our loggers were not placed until hatch. Ffith, in our sample, there was less temperature variability among nests with high feeding levels than those with medium or low feeding levels. Finally, it is possible that parental care behavior was affected by collection of morphological and physiological data before our observations, though we attempted to minimize these effects by waiting until birds appeared to resume normal behavior before beginning parental care data collection.

This study also has several strengths. In 113 wild nestling barn swallows, we investigated fine-scale heterogeneity in temperature associations with nestling mass before fledging across ontogeny and social environments, whereas most previous work has focused on characterizing overall patterns in a population or on documenting broader-scale environmental dependence (e.g., across climatic zones). We tracked temperature continuously at the nest-level, rather than at site- or region-level, as many previous studies have done. The fine-scale resolution of our temperature data enabled us to more precisely quantify variation in temperature associations with mass among individuals within the same population, allowing us to investigate the role of developmental timing and social environment in shaping temperature effects on nestlings. Additionally, we characterized average nest feeding levels using detailed measures of parental care behavior collected longitudinally across the nestling period, allowing us to thoroughly characterize average nest-level differences in feeding and assess whether parental feeding can buffer nestlings from the negative impacts of extreme and variable temperatures. In addition, by investigating three temperature variables (minimum temperature, maximum temperature, temperature variability) that capture different aspects of the thermal environment, our work offers insight on the effects hot, cold, and variable temperatures on nestlings. We provide a more complete picture of how different aspects of the thermal environment affect nestling growth, while broadening our understanding of how developmental and social factors may exacerbate or ameliorate the impacts of climate change on vulnerable individuals.

## Summary and Conclusions

In a study of wild barn swallow nestlings, we found that the associations of temperature with nestling mass differed among three temperature variables—lower minimum temperatures, higher maximum temperatures, and greater temperature variability were associated with lower nestling mass before fledging. Further, the associations of minimum temperature with nestling mass differed before versus after putative development of thermoregulatory independence. The associations of temperature with nestling mass did not vary with relative nestling size within a brood. Finally, we found evidence that the associations of extreme and variable temperature with nestling mass depended on level of parental feeding, with higher levels of parental feeding ameliorating the negative effects of temperature. Although prior work has revealed broad-scale environmental dependence of temperature effects on nestlings, our study unveils factors shaping fine-scale heterogeneity in temperature effects on nestlings. We offer initial insight into how early-life developmental constraints imposed by timing of thermoregulatory development, competition with brood mates, and amount of care provided by parents shape the impacts of extreme and variable temperatures on individual birds, providing the basis for future research to tease apart the mechanisms through which climate change affects development.

## Acknowledgements

We thank Aleea Pardue, Marina Ayala, Heather Kenny-Duddela, Avani Fachon, Grant Gonzalez, and Molly McDermott for assistance and/or advice regarding field work. We thank the private landowners who give us access to their property where barn swallows nest. Finally, we thank members of the Safran, Patricelli, and Karp labs for their constructive feedback on this project.

## Supporting information

**S1 Fig. A directed acyclic graph displaying hypothesized causal relationships investigated in question one: does the effect of temperature on nestling mass differ when the exposure is assessed during early versus late development?** The explanatory and response variables have black boxes, while confounders, precision covariates, and random intercepts have gray boxes. Below the DAG are the formulas for the linear mixed models we ran separately for each temperature variable (minimum, maximum, and variability).

**S2 Fig. A directed acyclic graph displaying hypothesized causal relationships investigated in question two: does the effect of temperature on nestling mass depend on whether a nestling is the smallest in the brood?** The explanatory and response variables have black boxes, while confounders, precision covariates, and random intercepts have gray boxes. Below the DAG is the formula for the linear mixed models we ran separately for each temperature variable (minimum, maximum, and variability).

**S3 Fig. A directed acyclic graph displaying hypothesized causal relationships investigated in question three: does the effect of temperature on nestling mass before fledging depend on the average amount of feeding provided by parents?** The explanatory and response variables have black boxes, while confounders, precision covariates, and random intercepts have gray boxes. Below the DAG is the formula for the linear mixed models we ran separately for each temperature variable (minimum, maximum, and variability).

**S4 Fig. Best unbiased linear predictions (BLUPs) for the feeding rate (visits/hour) at each nest across development in wild barn swallows in Boulder County, CO.** A boxplot of feeding BLUPs, measured in visits per hour, is provided for each of the three levels of parental care (‘low,’ ‘med,’ ‘high’) used for stratified analyses in question three.

**S5 Fig. Daily temperatures recorded at wild barn swallow nests in Boulder County, CO.** Daily minimum temperature (a), maximum temperature (b), and temperature variability (interquartile range) (c) in degrees Celsius recorded by Govee thermometers near each barn swallow nest during the nestling rearing period. Each color corresponds to one of seven breeding sites. Lines connect daily measures for each individual nest.

**S1 Table. Ethogram of barn swallow parental care behaviors.** Each documented behavior has a corresponding location, type (state versus event), description, and developmental context. This ethogram was used for parental care observations.

**S2 Table. Background characteristics of wild nestling barn swallows in Boulder County, CO.** Total number of feeding visits, nestling mass, and nestling wing length are separately estimated for three time points (days three to four, days eight to nine, and days 11-13). Temperature variability is defined as the interquartile range. The table provides the sample size for each variable at the level at which it was collected (n), as well as the mean, standard deviation (SD), minimum, and maximum.

**S3 Table. Associations of three temperature variables in early and late development (before or after six days post-hatch) with nestling mass for wild barns swallows in Boulder County, CO.** Temperature variability is defined as the interquartile range. For each temperature variable, results are provided for unadjusted and adjusted models. Sample size for each developmental stage is provided in the header (n). For each model, the table provides the effect size and 95% confidence interval for temperature effects (β (95% CI)), and the corresponding degrees of freedom, t-value, and p-value from a two-tailed t-test using a Satterthwaite degree of freedom estimation.

**S4 Table. Associations of three temperature variables with nestling mass, assessed in separate models stratified by relative nestling size at days eight to nine measure (smallest vs. other), for wild barn swallows in Boulder County, CO.** Temperature variability is defined as the interquartile range. For each temperature variable, results are provided for unadjusted and adjusted models. Sample size for each stratum is provided in the header (n). For each model, the table provides the effect size and 95% confidence interval for temperature effects (β (95% CI)), and the corresponding degrees of freedom, t-value, and p-value from a two-tailed t-test using a Satterthwaite degree of freedom estimation.

**S5 Table. Associations of three temperature variables with nestling mass, assessed in separate models stratified by levels of parental feeding, for wild barn swallows in Boulder County, CO.** Temperature variability is defined as the interquartile range. For each temperature variable, results are provided for unadjusted and adjusted models. Sample size for each stratum is provided in the header (n). For each model, the table provides the effect size and 95% confidence interval for temperature effects (β (95% CI)), and the corresponding degrees of freedom, t-value, and p-value from a two-tailed t-test using a Satterthwaite degree of freedom estimation.

