## Supplementary material for "The effects of temperature on nestling growth in a songbird depend on developmental and social context": S1 Fig

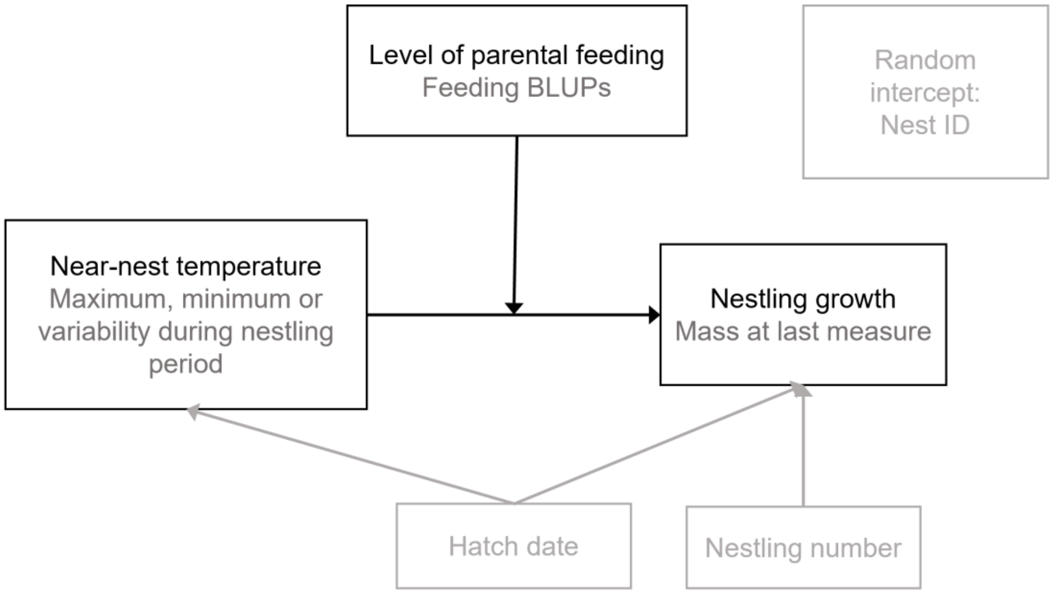


*One model with the following structure for each parental feeding strata (low, medium, high) and temperature variable (nine models total):*

Mass ~ temperature + nestling_number + hatch_date + (1|nest_id)

**S1 Fig. A directed acyclic graph displaying hypothesized causal relationships investigated in question one: does the effect of temperature on nestling mass differ when the exposure is assessed during early versus late development?** The explanatory and response variables have black boxes, while confounders, precision covariates, and random intercepts have gray boxes. Below the DAG are the formulas for the linear mixed models we ran separately for each temperature variable (minimum, maximum, and variability).
