## Supplementary material for "The effects of temperature on nestling growth in a songbird depend on developmental and social context": S1 Table

**S1 Table. Ethogram of barn swallow parental care behaviors.** Each documented behavior has a corresponding location, type (state versus event), description, and developmental context. This ethogram was used for parental care observations.

| Behavior location | Behavior name | Behavior type | Description | Developmental context |
|---|---|---|---|---|
| At nest | Feeding | Event | The parent delivers food as indicated by putting its beak into the mouth of the nestling. Prey often cannot be seen in the parent's bill. Feeding usually occurs immediately after arrival at the nest. | Day 3: nestlings are small, so they usually cannot be seen by the observer. Feeding visits are identified by the parent leaning into the nest immediately after arrival from foraging |
| | | | | Day 8: the parent can often be seen inserting its bill into the gap of nestlings. The parent usually still leans into the nest slightly when feeding. |
| | | | | Day 12: nestlings can be seen and feeding occurs without leaning into the nest. |
| | Brooding | State | The parent settles itself all the way into the nest on top of the nestlings. The parent may turn around in the nest while brooding. Brooding behavior looks similar across nestling ages but is far more common with younger nestlings (Day 3). | NA |
| | Sanitizing | State | The parent removes nestling fecal sacs or rearranges nestlings and feathers in the nest. The parent may carry fecal sacs out of the nest or consume them. When rearranging | Day 3: The parents often sanitize while settled in the nest cup. The parent will shift around and the tail will twitch. The head will duck down into the nest. |

| | | | | |
|---|---|---|---|---|
| | | | feathers or nestlings, the parent may perch on the edge of the nest and lean in deeply, tail twitching (see-saw motion; head and tail pivot up and down). The parent may also sanitize while brooding, as indicated by their head in the nest and tail twitching. | Days 8 and 12: sanitizing more common occurs while perched on the nest edge. Parents may be seen carrying fecal sacks away. |
| | Preening | State | The parent uses the beak to clean, stroke, or re-position its feather while perched on the nest rim or settled in the nest brooding. | NA |
| | Perching | State | The parent stands with feet on the side of the nest, not engaged in any other behaviors. The parent may look around the barn or it may peer inside the nest. | NA |
| Away from nest | Near nest | State | The parent is visible at their typical perch location(s). Importantly here it is reasonable to assume that the bird can see the nest (i.e. they are able to be vigilant of any nest threats). | NA |
| | Far from nest | State | The parent flies away from the nest and is either observed leaving the barn or headed towards a barn opening but that might be out of sight for the observer. | NA |
| | Unknown | State | It is not known if the parent can reasonably see the nest or is in the barn at all. | NA |
