## Supplementary material for "The effects of temperature on nestling growth in a songbird depend on developmental and social context": S2 Fig

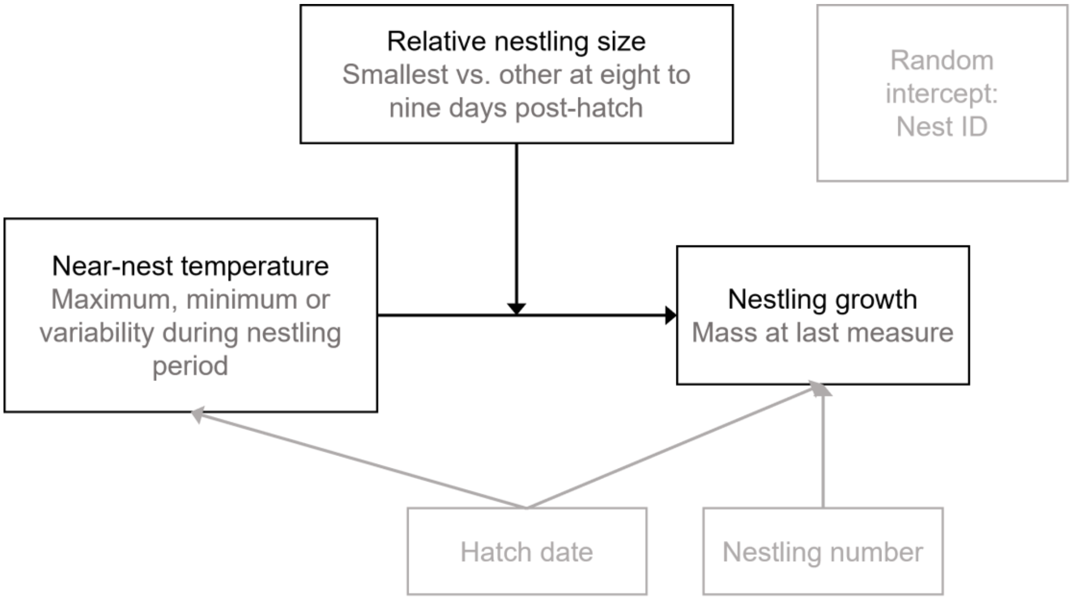


*One model with the following structure for each relative nestling size strata (smallest, other) and temperature variable (six models total):*

Mass ~ temperature + nestling_number + hatch_date + (1|nest_id)

**S2 Fig. A directed acyclic graph displaying hypothesized causal relationships investigated in question two: does the effect of temperature on nestling mass depend on whether a nestling is the smallest in the brood?** The explanatory and response variables have black boxes, while confounders, precision covariates, and random intercepts have gray boxes. Below the DAG is the formula for the linear mixed models we ran separately for each temperature variable (minimum, maximum, and variability).
