## Supplementary material for "The effects of temperature on nestling growth in a songbird depend on developmental and social context": S2 Table

**S2 Table. Background characteristics of wild nestling barn swallows in Boulder County, CO.** Total number of feeding visits, nestling mass, and nestling wing length are separately estimated for three time points (days three to four, days eight to nine, and days 11-13). Temperature variability is defined as the interquartile range. The table provides the sample size for each variable at the level at which it was collected (n), as well as the mean, standard deviation (SD), minimum, and maximum.

| Variable | n | Mean | SD | Minimum | Maximum |
|---|---|---|---|---|---|
| **Near-nest temperature** | | | | | |
| Minimum temperature (C) | 31 | 13.10 | 1.80 | 10.30 | 17.10 |
| Maximum temperature (C) | 31 | 36.89 | 2.96 | 29.10 | 42.30 |
| Temperature variability (C) | 31 | 9.26 | 2.43 | 5.35 | 13.60 |
| **Other nest characteristics** | | | | | |
| Brood size (number of nestlings) | 31 | 3.42 | 0.96 | 1.00 | 5.00 |
| Hatch date (days since June 1) | 31 | 20.58 | 10.05 | 2.00 | 38.00 |
| Nest age at last measure (days since hatch) | 31 | 12.26 | 0.51 | 11.00 | 13.00 |
| **Total feeding rate (visits/hour) across development**[1] | | | | | |
| Days three to four | 30 | 12.21 | 6.92 | 3.00 | 35.73 |
| Days eight to nine | 28 | 12.73 | 8.08 | 2.97 | 29.92 |
| Days 11-13 | 30 | 21.62 | 12.94 | 3.98 | 60.35 |
| **Right wing length (mm) across development** | | | | | |
| Days three to four | 113 | 9.40 | 1.94 | 5.00 | 13.50 |
| Days eight to nine | 108 | 30.71 | 5.36 | 15.00 | 40.50 |
| Days 11-13 | 106 | 52.78 | 6.44 | 31.00 | 67.00 |
| **Nestling mass (g) across development** | | | | | |
| Days three to four | 113 | 4.00 | 1.30 | 1.25 | 7.73 |
| Days eight to nine | 108 | 12.95 | 2.56 | 6.01 | 17.66 |
| Days 11-13 | 106 | 17.57 | 2.21 | 10.35 | 22.36 |

[1]Parental feeding behaviors are measured at the level of the nest and include the totals for both social parents (maternal and paternal care).
