## Supplementary material for "The effects of temperature on nestling growth in a songbird depend on developmental and social context": S3 Fig

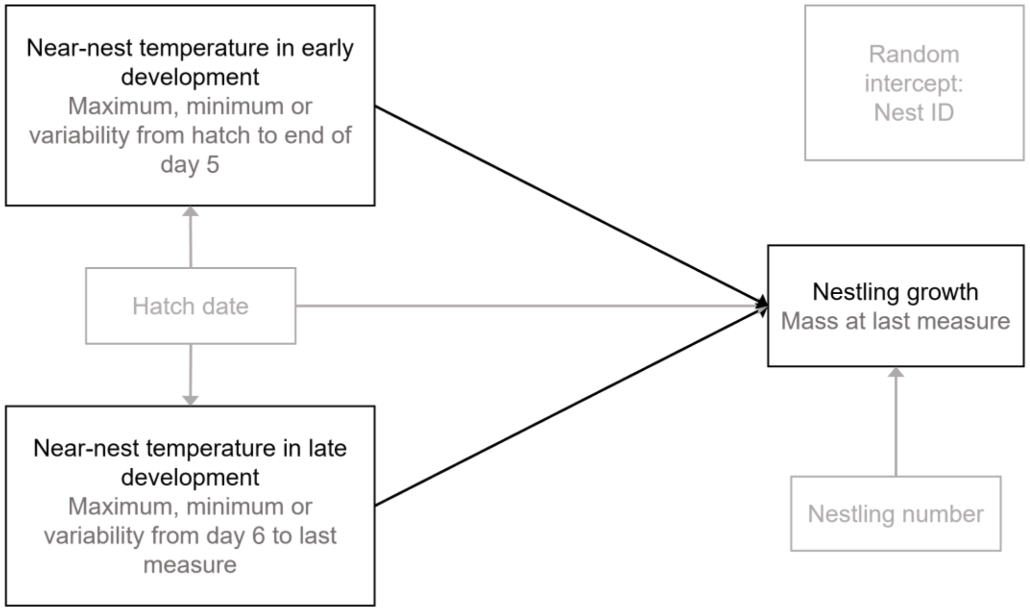


*One of each of the following model structures for each temperature variable (six models total):*

Mass ~ before_temperature + nestling_number + hatch_date + (1|nest_id)

Mass ~ after_temperature + nestling_number + hatch_date + (1|nest_id)

**S3 Fig. A directed acyclic graph [3] displaying hypothesized causal relationships investigated in question three: does the effect of temperature on nestling mass before fledging depend on the average amount of feeding provided by parents?** The explanatory and response variables have black boxes, while confounders, precision covariates, and random intercepts have gray boxes. Below the DAG is the formula for the linear mixed models we ran separately for each temperature variable (minimum, maximum, and variability).
