## Supplementary material for "The effects of temperature on nestling growth in a songbird depend on developmental and social context": S3 Table

**S3 Table. Associations of three temperature variables in early and late development (before or after six days post-hatch) with nestling mass for wild barns swallows in Boulder County, CO.** Temperature variability is defined as the interquartile range. For each temperature variable, results are provided for unadjusted and adjusted models. Sample size for each developmental stage is provided in the header (n). For each model, the table provides the effect size and 95% confidence interval for temperature effects (β (95% CI)), and the corresponding degrees of freedom, t-value, and p-value from a two-tailed t-test using a Satterthwaite degree of freedom estimation.

| | Before models (n = 106) | | | | After models (n = 106) | | | |
|---|---|---|---|---|---|---|---|---|
| Type | β (95% CI)[1] | DF | t-value | p-value | β (95% CI)[1] | DF | t-value | p-value |
| **Effect of minimum temperature** | | | | | | | | |
| Unadjusted | 1.11 (0.45, 1.80) | 28.58 | 3.22 | 0.003 | 0.50 (-0.22, 1.25) | 28.55 | 1.30 | 0.20 |
| Adjusted[2] | 1.16 (0.32, 2.00) | 26.96 | 2.79 | 0.01 | 0.29 (-0.45, 1.06) | 26.96 | 0.75 | 0.46 |
| **Effect of maximum temperature** | | | | | | | | |
| Unadjusted | -0.87 (-1.53, -0.24) | 29.13 | -2.61 | 0.01 | -1.16 (-1.82, -0.53) | 28.51 | -3.56 | 0.001 |
| Adjusted[3] | -0.66 (-1.32, -0.04) | 27.67 | -2.03 | 0.05 | -1.13 (-1.76, -0.52) | 26.87 | -3.60 | 0.001 |
| **Effect of temperature variability** | | | | | | | | |
| Unadjusted | -1.32 (-1.9, -0.78) | 28.61 | -4.53 | 0.0001 | -1.39 (-1.99, -0.85) | 28.45 | -4.74 | 0.0001 |
| Adjusted[4] | -1.41 (-2.07, -0.77) | 27.00 | -4.21 | 0.0003 | -1.33 (-1.95, -0.70) | 26.38 | -4.14 | 0.0003 |

[1]Estimated β (95% CI) from stratified linear mixed models in which temperature before or after thermoregulatory independence are the explanatory variables of interest, nestling mass is the outcome of interest, and nest ID was included as a random intercept. Adjusted models include hatch date and number of nestlings in the nest. Continuous predictors as z-score standardized.

[2]R-squared for adjusted minimum temperature models. Before model: Marginal R-squared = 0.23, Conditional R-squared = 0.82; After model: Marginal R-squared = 0.05, Conditional R-squared = 0.82

[3]R-squared for adjusted maximum temperature models. Before model: Marginal R-squared = 0.14, Conditional R-squared = 0.81; After model: Marginal R-squared = 0.25, Conditional R-squared = 0.82

[4]R-squared for adjusted temperature variability models. Before model: Marginal R-squared = 0.34, Conditional R-squared = 0.81; After model: Marginal R-squared = 0.37, Conditional R-squared = 0.81
