## Supplementary material for "The effects of temperature on nestling growth in a songbird depend on developmental and social context": S4 Fig

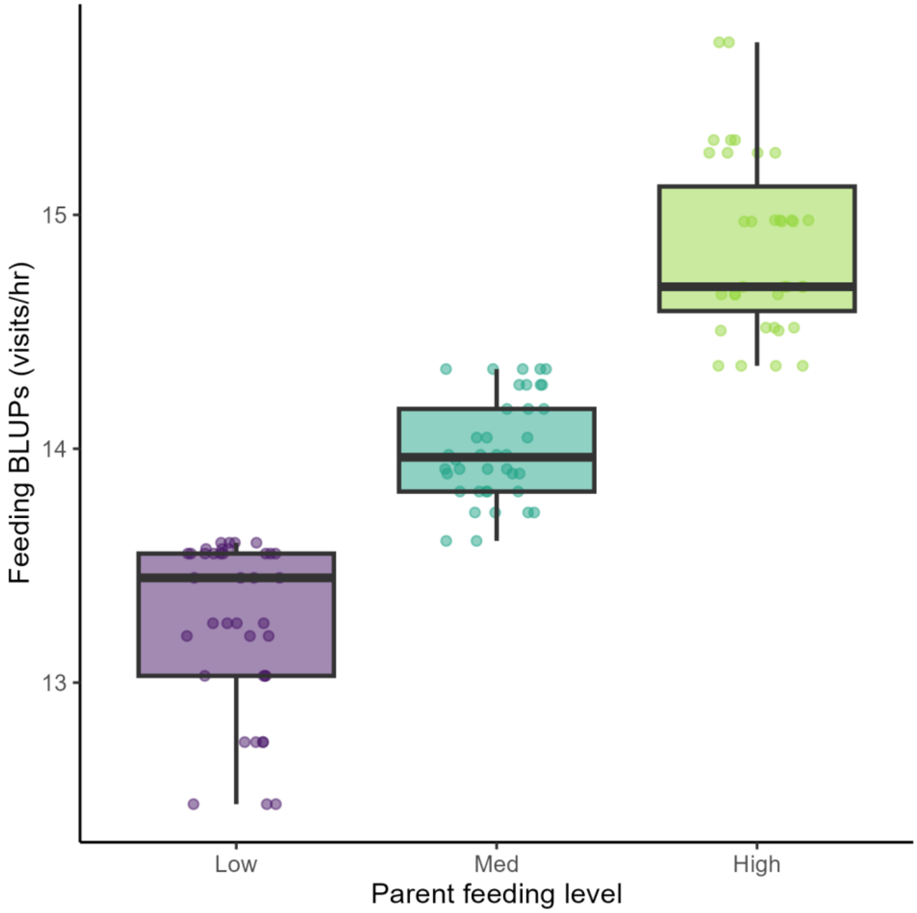


**S4 Fig. Best unbiased linear predictions (BLUPs) for the feeding rate (visits/hour) at each nest across development in wild barn swallows in Boulder County, CO.** A boxplot of feeding BLUPs, measured in visits per hour, is provided for each of the three levels of parental care (‘low,’ ‘med,’ ‘high’) used for stratified analyses in question three.
