## Supplementary material for "The effects of temperature on nestling growth in a songbird depend on developmental and social context": S5 Fig

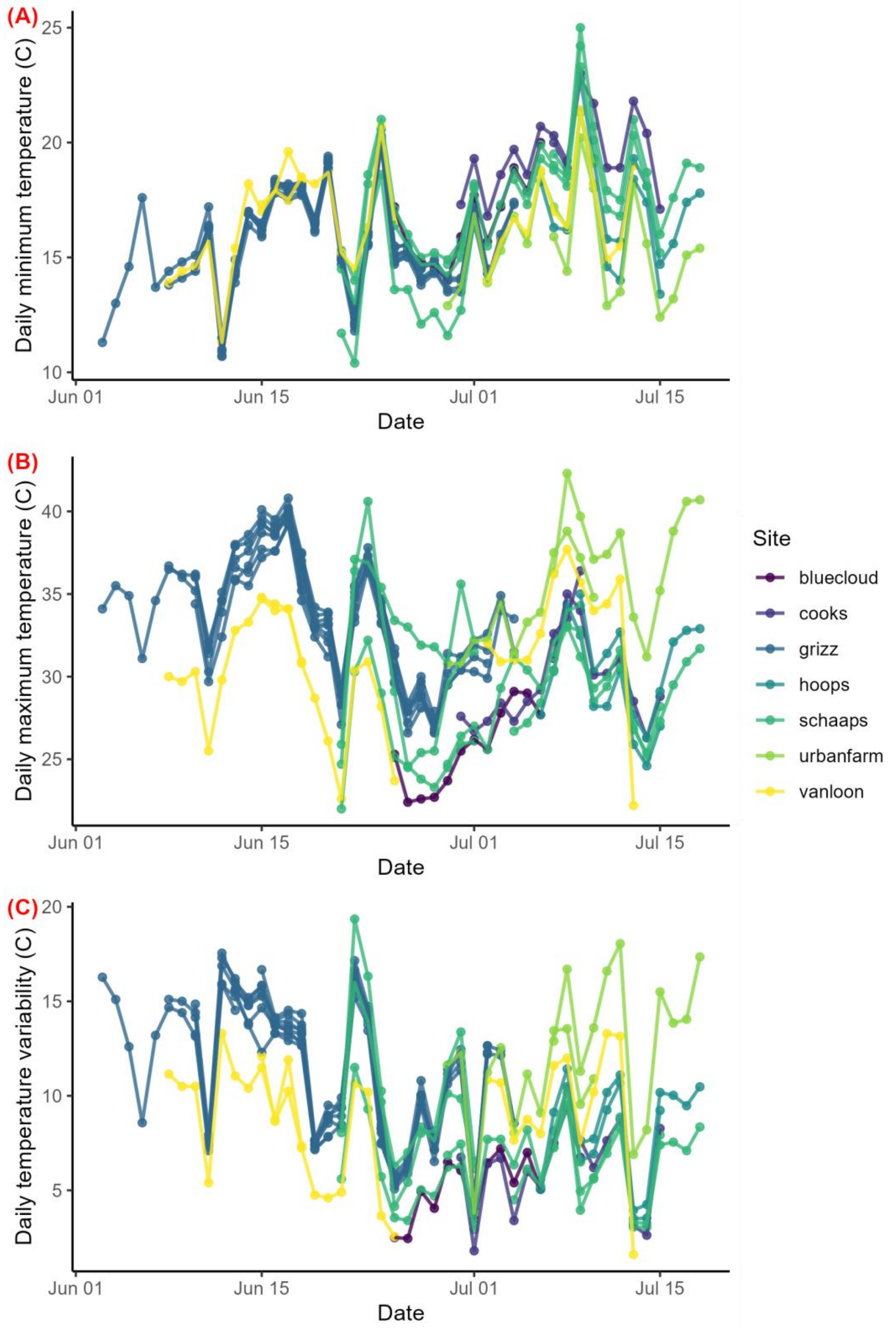


**S5 Fig. Daily temperatures recorded at wild barn swallow nests in Boulder County, CO.** Daily minimum temperature (**a**), maximum temperature (**b**), and temperature variability (interquartile range) (**c**) in degrees Celsius recorded by Govee thermometers near each barn swallow nest during the nestling rearing period. Each color corresponds to one of seven breeding sites. Lines connect daily measures for each individual nest.
