## Supplementary material for "The effects of temperature on nestling growth in a songbird depend on developmental and social context": S5 Table

**S5 Table. Associations of three temperature variables with nestling mass, assessed in separate models stratified by levels of parental feeding, for wild barn swallows in Boulder County, CO.** Temperature variability is defined as the interquartile range. For each temperature variable, results are provided for unadjusted and adjusted models. Sample size for each stratum is provided in the header (n). For each model, the table provides the effect size and 95% confidence interval for temperature effects (β (95% CI)), and the corresponding degrees of freedom, t-value, and p-value from a two-tailed t-test using a Satterthwaite degree of freedom estimation.

| | Low parental feeding models (n = 35) | | | | Medium parental feeding models (n = 36) | | | | High parental feeding models (n = 35) | | | |
|---|---|---|---|---|---|---|---|---|---|---|---|---|
| Type | β (95% CI)[1] | DF | t-value | p-value | β (95% CI)[1] | DF | t-value | p-value | β (95% CI)[1] | DF | t-value | p-value |
| **Effect of minimum temperature** | | | | | | | | | | | | |
| Unadjusted | 0.96 (-0.09, 1.99) | 7.98 | 1.77 | 0.12 | 1.89 (0.68, 3.04) | 9.13 | 3.23 | 0.01 | -0.20 (-1.03, 0.60) | 7.03 | -0.49 | 0.64 |
| Adjusted[2] | 1.73 (-0.11, 3.53) | 5.99 | 1.89 | 0.11 | 2.36 (0.68, 3.92) | 6.89 | 2.70 | 0.03 | -0.53 (-2.05, 0.94) | 4.90 | -0.70 | 0.52 |
| **Effect of maximum temperature** | | | | | | | | | | | | |
| Unadjusted | -1.06 (-2.01, -0.12) | 8.62 | -2.24 | 0.05 | -1.65 (-2.86, -0.45) | 8.90 | -2.72 | 0.02 | 0.19 (-0.54, 0.91) | 8.35 | 0.52 | 0.62 |
| Adjusted[3] | -1.08 (-2.44, 0.34) | 6.18 | -1.62 | 0.16 | -1.22 (-2.34, -0.04) | 7.40 | -2.11 | 0.07 | 0.01 (0.91, 0.92) | 5.61 | 0.03 | 0.98 |
| **Effect of temperature variability** | | | | | | | | | | | | |
| Unadjusted | -1.11 (-2.05, -0.13) | 8.24 | -2.27 | 0.05 | -2.06 (-2.97, -1.15) | 8.76 | -4.44 | 0.002 | 0.08 (-0.70, 0.89) | 7.53 | 0.21 | 0.84 |
| Adjusted[4] | -1.35 (-2.77, 0.14) | 6.07 | -1.88 | 0.11 | -1.8 (-2.78, -0.75) | 7.42 | -3.46 | 0.01 | -0.01 (-1.25, 1.28) | 5.64 | -0.01 | 0.99 |

[1]Estimated β (95% CI) from stratified linear mixed models in which temperature is the explanatory variable of interest, nestling mass is the outcome of interest, and nest ID was included as a random intercept. Adjusted models include hatch date and number of nestlings in the nest. Continuous predictors are z-score standardized.

[2]R-squared for adjusted minimum temperature models. Low parental feeding model: Marginal R-squared = 0.3, Conditional R-squared = 0.87; Medium parental feeding model: Marginal R-squared = 0.52, Conditional R-squared = 0.83; High parental feeding model: Marginal R-squared = 0.1, Conditional R-squared = 0.62

[3]R-squared for adjusted maximum temperature models. Low parental feeding model: Marginal R-squared = 0.2, Conditional R-squared = 0.87; Medium parental feeding model: Marginal R-squared = 0.46, Conditional R-squared = 0.84; High parental feeding model: Marginal R-squared = 0.07, Conditional R-squared = 0.63

[4]R-squared for adjusted temperature variability models. Low parental feeding model: Marginal R-squared = 0.27, Conditional R-squared = 0.87; Medium parental feeding model: Marginal R-squared = 0.6, Conditional R-squared = 0.82; High parental feeding model: Marginal R-squared = 0.07, Conditional R-squared = 0.63
